# Family-Companion: analyse, visualise, browse, query and share your homology clusters

**DOI:** 10.1101/266742

**Authors:** Ludovic Cottret, Martial Briand, Corinne Rancurel, Sébastien Carrere

## Abstract

Identifying homology groups in predicted proteomes from different biological sources allows biologists to address questions as diverse as inferring species-specific proteins or retracing the phylogeny of gene families. Nowadays, command-line software exists to infer homology clusters. However, computing and interpreting homology groups with this software remains challenging for biologists and requires computational skills.

We propose Family-Companion, a web server dedicated to the computation, the analysis and the exploration of homology clusters. Family-Companion aims to fill the gap between analytic software and databases presenting orthologous groups based on a set of public data. It offers a user-friendly interface to launch or upload precomputed homology cluster analysis, to explore and share the results with other users. The exploration of the results is highly facilitated by interactive solutions to visualize proteome intersections via Venn diagrams, phylogenetic trees, multiple alignments, and also by querying the results by blast or by keywords.

Family-Companion is available at http://family-companion.toulouse.inra.fr with a demo dataset and a set of video tutorials. Source code and installation protocol can be found at https://framagit.org/BBRIC/family-companion/. A container-based package simplifies the installation of the web-suite.

## 1. Introduction

The proteome of an organism is the set of proteins encoded by its genome. Proteins that derive from a common ancestral protein are called homologous proteins. A common assumption is to consider that orthologous proteins (homologous proteins that diverged after a speciation event) have a similar biological role in different species. In contrast, paralogous proteins are homologous proteins that have diverged after a duplication event within one species. By comparing all protein sequences found in a set of organisms, it is possible to classify them in homology (orthology/paralogy) clusters. These can then be analysed to identify, for instance, proteins specific to a group of species that share a biological trait of interest (e.g. pathogenicity or symbiosis (1, 2). Besides, identification of universally conserved single copy proteins can help to infer species phylogeny (3). Finally, analysis of presence/absence and copy number of proteins in different homology groups across species allows their evolutionary history to be inferred (4).

The identification of homology groups from proteomes can be performed thanks to command-line software based on sequence comparison and clustering (5–7). However, the results, in the form of complex and non standardized text files, are hardly exploitable or interpretable without computational skills. Existing web resources provide solutions to facilitate the exploration of pre-computed homology groups. Orthomcl-db (8) and OrthoDB (9) allow complex queries to be made from several characteristics such as the inclusion or exclusion of taxa, statistics or functional annotations. EggNogg (10) proposes an online visualization of trees that show duplication and speciation events to distinguish between paralogs and orthologs. Orthomcl-db, OrthoDB and EggNogg all propose a catalogue of orthologs spanning a large scope of species. They also allow user-provided proteomes to be mapped onto pre-computed homology groups.

However, none of these web servers provide a combined interface to calculate homology groups from a user-defined set of proteomes of interest, and to explore them through online queries and visualisation tools. OrthoVenn (11) provides tools for cluster annotation and Venn diagrams from user-provided protein sequences, even if the number of proteomes is limited to 6. Spocs (12) provides a graph-based ortholog prediction method and tools to visualise predicted ortholog/paralog relationships.

In order to supplement existing offers, we propose Family-Companion (mentioned below as F-C), a rich web application that allows computation, exploration and sharing of homology groups.

## 2. Results

An overview of the analysis flow in F-C is illustrated in Supp Fig 1.

### 2.1. Homology cluster computation and automatic analyses

#### Inference of homology clusters

A dynamic online form enables users to upload an unlimited number of proteomes in FASTA format. Once all the proteomes of interest have been uploaded, homology groups can be inferred by automatically launching OrthoMCL with parameters that can be easily tuned (Supp. Fig. 2 and Supp. Fig. 3). If the user has already generated homology groups using OrthoMCL v1.4 or V 2, Orthofinder Synergy or any other tool generating a compatible format, results can be directly uploaded to Family Companion for exploration and visualization (Supp. Fig. 2).

#### Alignments and phylogenetic trees

For each homology group, a multi-alignment and a phylogenetic tree are computed to facilitate the analysis of protein families.

#### Generation of core, pan proteomes and species-specific proteins

The core-proteome corresponds to single-copy proteins conserved in all analysed proteomes. This set of one-to-one conserved orthologous proteins can be particularly useful to infer species phylogenies. To facilitate their reconstruction, F-C provides a super alignment built by concatening alignments of proteins involved in each one-to-one ortholog cluster. Of note, the content of the core proteome depends highly on the parameters used for protein clustering. The pan-proteome provides a representation of the diversity of proteins present in the set of organisms of interest. It is composed of one representative of each homology group and all the species-specific single copy proteins. The user can optionally tag a ‘reference’ proteome. In this case the proteins from the reference proteome will be selected as representative in each homology group in which the species is present. Otherwise, a matrix is built by computing a distance between each sequence. The sequence with the minimal sum of distances is chosen as representative.

For each proteome, a set of specific proteins is computed. It is composed of single-copy proteome-specific proteins (i.e. all the proteins that are not present in any of the homology groups) and multi-copy proteome-specific proteins (i.e. proteins in homology groups present in a unique species).

The core-proteome super-alignment, the pan-proteome and the specific proteins are all downloadable in FASTA format (Supp. Fig. 4).

#### Generation of phylogenetic profiles

In order to assess the presence, expansion or reduction of protein families across species, F-C computes presence/absence and abundance matrices. In these downloadable matrices, rows correspond to the homology groups, and columns to taxa. The cells either include a Boolean value (presence / absence) or copy numbers of proteins belonging to a homology group in the corresponding taxon.

#### Functional annotation of homology clusters

InterPro annotations (13) can be uploaded in the main F-C online form (Supp. Fig. 3). Then, functional annotations are assigned to homology groups by concatenating the annotations of the proteins composing them.

### 2.2 Interactive visualization solutions

### Alignments and phylogenetic trees

For each homology group, a multi-alignment and a phylogenetic tree are displayed in an interactive way (Supp. Fig. 5) This facilitates the analysis of the evolutionary events (speciation, duplication) that link the proteins of the homology group.

### Visualization of phylogenetic profiles

Each phylogenetic profile can be visualized online with an interactive heatmap (Supp. Fig. 6) where clustering trees for rows (homology groups) and columns (proteomes) are displayed.

### Interactive charts and Venn diagrams

A series of interactive charts is produced at each analysis to visualize differences between the selected proteomes. For instance, one can visualize the number of proteins classified into homology groups, or specific to particular taxa (Supp. Fig. 7).

F-C provides interactive Venn diagrams to compare proteomes based on their content in homology groups (Supp. Fig. 8). By clicking on diagram intersections, links to corresponding homology clusters are displayed.

### 2.3 Browse and Query

The web interface provides an immediate navigation between tables listing proteins, homology groups, alignments, trees and phylogenetic profiles (Supp. Fig. 9). In order to explore results of homology groups in more detail, F-C provides a Query Builder. A search engine returns the groups to which a particular protein belongs via its accession number or a keyword present in its annotation. An integrated Blast (14) server can also find homology clusters through sequence similarity to a query protein (Supp. Fig. 10).

Moreover, a form allows queries to be graphically constructed to extract the homology clusters present in a set of proteomes and absent in another set (Supp. Fig. 11). This can be used to investigate clade-specific functions.

### 2.4 Share private analyses

By default, new analyses are private and only accessible by the authenticated user who uploaded data. However, this user has the possibility to generate a public URL in order to make the results available to others. This stable URL allows the link to be integrated into analyses in other pages or simply to share them with collaborators. The sharing of an analysis can also be cancelled by a simple click (Supp. Fig. 12).

## 3. Implementation

The back-end part is implemented in Perl. The front-end part is implemented in Javascript. The framework Ext Js 6.0 [^1^ https://www.sencha.com/products/extjs] was used for the user interface. Interactive charts use the HighCharts library [^2^ https://www.highcharts.com]. Venn diagrams are displayed thanks to the Jvenn library (15). Interactive heatmaps are built with the Inchlib JavaScript and python libraries (16). Alignments are displayed with the BioJs MSA viewer library (17). Phylogenetic trees are visualised thanks to the Phylocanvas javascript library [^3^ http://phylocanvas.org/]. New homology clusters are inferred thanks to a modified version of OrthoMcl V1.4 (5) using Diamond (18) and Parallel (19) software to speed up similarity computing between sequences. Multiple alignments are computed with Mafft (20) and cleaned with Trimal (21). Phylogenetic trees are inferred by FastTree (22).

## 4. Conclusion

The user-friendly interface of F-C provides powerful, interactive web solutions to explore homology clusters and address the most common questions in this kind of approach. Another interesting feature is to handle homology clusters for a set of proteomes provided by the users themselves, which complements the existing solutions. In addition, sharing private analyses facilitates collaborative work.

Future improvements can be imagined. Annotation transfer could be facilitated between the proteomes. Statistical analyses such as Principal Component Analysis could be automatically performed to describe proteomes according to the homology clusters. Finally, an incremental update of clusters and analyses could save time when new proteomes are provided.

## Acknowledgements

We are grateful to the genotoul bioinformatics platform Toulouse Midi-Pyrenees (Bioinfo Genotoul) for hosting F-C. We thank Etienne Danchin, Matthieu Barret and Jérôme Gouzy for testing F-C, reading the manuscript and providing helpful comments and Clare Gough for checking language mistakes.

